# Developmental regulation and functional prediction of microRNAs in an expanded *Fasciola hepatica* miRNome

**DOI:** 10.1101/2021.11.08.467689

**Authors:** Caoimhe M. Herron, Anna O’Connor, Emily Robb, Erin McCammick, Claire Hill, Nikki J. Marks, Mark W. Robinson, Aaron G. Maule, Paul McVeigh

## Abstract

1

The liver fluke, *Fasciola hepatica*, is a global burden on the wellbeing and productivity of farmed ruminants, and a zoonotic threat to human health. Despite the clear need for accelerated discovery of new drug and vaccine treatments for this pathogen, we still have a relatively limited understanding of liver fluke biology and host interactions. Noncoding RNAs, including micro (mi)RNAs, are key to transcriptional regulation in all eukaryotes, such that an understanding of miRNA biology can shed light on organismal function at a systems level. Four previous publications have reported up to 89 mature miRNA sequences from *F. hepatica*, but our data show that this does not represent a full account of this species miRNome. We have expanded on previous studies by sequencing, for the first time, miRNAs from multiple life stages (adult, newly excysted juvenile (NEJ), metacercariae and adult-derived extracellular vesicles (EVs)). These experiments detected an additional 61 high-confidence miRNAs, most of which have not been described in any other species, expanding the *F. hepatica* miRNome to 150 mature sequences. We used quantitative (q)PCR assays to provide the first developmental profile of miRNA expression across metacercariae, NEJ, adult and adult-derived Evs. The majority of miRNAs were expressed most highly in metacercariae, with at least six distinct expression clusters apparent across life stages. Intracellular miRNAs were functionally analysed to identify target mRNAs with inversely correlated expression in *F. hepatica* tissue transcriptomes, highlighting regulatory interactions with key virulence transcripts including cathepsin proteases, and neuromuscular genes that control parasite growth, development and motility. We also linked 28 adult-derived EV miRNAs with downregulation of 397 host genes in *F. hepatica*-infected transcriptomes from ruminant lymph node, peripheral blood mononuclear cell (PBMC) and liver tissue transcriptomes. These included genes involved in signal transduction, immune and metabolic pathways, adding to the evidence for miRNA-based immunosuppression during fasciolosis. These data expand our understanding of the *F. hepatica* miRNome, provide the first data on developmental miRNA regulation in this species, and provide a set of testable hypotheses for functional genomics interrogations of liver fluke miRNA biology.

**CONTRIBUTION TO THE FIELD:** Previous studies identified 89 micro (mi)RNAs (non-coding RNAs responsible for regulating gene expression) in the trematode parasite, *Fasciola hepatica*, and attached functional annotations to the miRNAs that are secreted by liver fluke *in vitro*. This study expands the known miRNA complement of *F. hepatica* by 40%, adding an additional 61 miRNAs, many of which appear *Fasciola*-specific. We used this expanded dataset to perform the first analysis of developmental miRNA expression across intra-mammalian parasites, showing clear developmentally regulated expression profiles across infectious metacercariae, juvenile and adult parasites, and extracellular vesicles secreted by adult parasites. We performed rigorous functional annotation of cellular miRNAs, correlating their expression with mRNA transcriptomes to identify potential roles for specific miRNAs in regulating expression of important proteases and nerve/muscle transcripts, all of which are important for parasite virulence. We also functionally analysed miRNAs secreted by adult parasites in terms of interactions with host transcripts; for the first time this analysis was supported by host transcriptome data, linking secreted miRNAs with mRNAs that are downregulated by fluke infection in sheep and cattle. This work contributes new data, insights and analyses to the field, providing a rich source of hypotheses for future experimental validation.

## 2 INTRODUCTION

*Fasciola* spp. liver fluke are important flatworm parasites of ruminants, impacting agricultural productivity and animal welfare worldwide, with *F. hepatica* and *F. gigantica* found predominantly in temperate and tropical regions respectively. Efforts to control these parasites rely almost entirely on blanket application of flukicidal anthelmintic drugs to ruminant herds, an approach which has resulted in a selective pressure favouring parasite populations carrying resistance alleles for most of the currently available flukicides (Kelley et al., 2016; Fairweather et al., 2020). Given the absence of a vaccine, new control options are urgently needed.

Understanding parasite biology and their host interactions is key to identifying new molecular targets for anthelmintic drugs, vaccines and non-chemical control options, and new biomarkers for improved diagnostics. Expanded availability of omics datasets have accelerated this process for *F. hepatica* (Cwiklinski and Dalton, 2018; McVeigh et al., 2018; McVeigh, 2020), but this species arguably still lags other parasites in the exploitation of these datasets for parasite control. One area where we fall short is the understanding of non-coding RNAs and post-transcriptional regulation of gene expression. Micro (mi)RNAs are non-coding RNAs responsible for the modulation of gene expression through RNA interference (RNAi) pathways. Most miRNAs bind to 3-prime untranslated region (3’ UTR) of their target transcripts, reducing mRNA stability and leading to downregulation of protein expression (O’Brien et al., 2018). This fundamental process occurs throughout eukaryotes, but our ability to probe and understand it in liver fluke specifically has been hindered by the lack of appropriate functional genomic protocols and an incomplete understanding of the miRNome. To date, 89 *F. hepatica* miRNAs have been reported in adult and newly excysted juvenile (NEJ) parasites (Xu et al., 2012b; Fontenla et al., 2015a; Fromm et al., 2015b; Ovchinnikov et al., 2020; Ricafrente et al., 2020). Given that miRBase (release 22.1) reports more than 100 miRNAs in other flatworm species, this is probably not the full extent of the *F. hepatica* miRNome. In addition, there are no published quantitative data on the developmental regulation of miRNA expression across *F. hepatica* life stages, insights which are essential in attaching function to these sequences. A recent study on *F. gigantica* (Hu et al., 2021) measured miRNA expression across intra-mammalian and intra-molluscan life stages, linking these sequences with regulation of metabolism, transport, growth and development. Given the miRNA homology shared between *F. hepatica* and *F. gigantica* (Hu et al., 2021) this dataset is a useful comparative tool that could yield insights for *F. hepatica* as well.

*F. hepatica* secretes miRNAs, which are of interest given their potential role in host-parasite interactions and as diagnostic biomarkers. Following the first report of extracellular vesicle (EV) release from *F. hepatica* (Marcilla et al., 2012), miRNAs were soon observed amongst the cargo molecules of these membrane bound carriers (Fromm et al., 2015b). These miRNAs have been analysed in terms of their potential to target immune transcripts of host organisms (Fromm et al., 2015b; Fromm et al., 2017; Ovchinnikov et al., 2020; Ricafrente et al., 2020), with data now showing that fhe-mir-125b can enter host immune cells and attach to host argonaut protein, forming a potentially functional silencing unit (Tran et al., 2021). Additional functional insight has come from using miRNA target prediction algorithms to identify the host mRNAs that are targeted for silencing by fluke secreted miRNAs. Ovchinnikov *et al*. focused on adult EV miRNAs, uding PITA and TargetSCAN to link 24 miRNAs to 321 cow and human mRNAs, including targets within the Wnt signalling pathway and the immune system (Ovchinnikov et al., 2020). Similarly, (Ricafrente et al., 2020) used miRTarget to link 38 NEJ expressed miRNAs (Fontenla et al., 2015a) with 26 target genes expressed in innate immune cells.

Our work builds on previous studies to catalogue new miRNAs within an expanded *F. hepatica* miRNome, alongside a developmental expression profile of these in muiltiple intra-mammalian life stages. We provide functional insight through rigorous miRNA target prediction analysis against both endogenous fluke tissue transcriptomes, and transcriptomes from sheep/cow tissues from fasciolosis infections. Network and co-expression analyses are used to categorise functions for both cellular and secreted miRNAs. This work significantly expands our understanding of the *F. hepatica* miRNome and provides a foundation for future functional genomics investigations into liver fluke miRNA biology.

## 3 MATERIALS AND METHODS

### 3.1 Parasite handling and sample preparation

Italian strain *F. hepatica* metacercariae were obtained from Ridgeway Research Ltd. These were either used directly for RNA extraction, or excysted to newly excysted juveniles (NEJs) as described previously (McVeigh et al., 2014). For generation of small RNA-Seq libraries, NEJs were maintained *in vitro* for 7 days either in RPMI 1640 (non-growing NEJs) or in RPMI 1640 supplemented with 20 % Foetal Bovine Serum (FBS) (growing NEJs). In later experiments for quantitative (q)PCR-based detection of miRNAs, NEJs were processed for RNA extraction within 3 hours of excystment. Comparisons between growing and non-growing samples were not performed by qPCR due to low availability of metacercariae at the time of experimentation. Adult parasites were recovered from sheep at abattoir (ABP Meats, Lurgan, Co Armagh, Northern Ireland) and snap frozen within 4h of collection. EVs were collected following incubation of adult parasites (5h, two flukes per ml), or NEJs (24h, 200 parasites per 500ul) in RPMI 1640, using ultracentrifugation as described previously (Cwiklinski et al., 2015b).

### 3.2 Small RNA sequencing and bioinformatics

In all cases, total RNA was extracted using Trizol Reagent (Thermo Fisher Scientific). For RNA-Seq experiments, RNA was extracted from 2000 metacercariae or newly excysted juvenile (NEJ) worms, from a single adult parasite, or from EV samples prepared as described above. RNA-Seq libraries were prepared using a TruSeq Small RNA Library Preparation Kit (Illumina), and sequenced, by Genome Quebec (McGill University Genome Quebec Innovation Centre, Montreal, Canada), on an Illumina Hi-Seq platform. Sequencing reads were adaptor trimmed with Cutdapt 2.5, and then processed for miRNA prediction using miRDeep2 (Friedlander et al., 2012). Reads were mapped against the WBPS13 version of the *F. hepatica* genome (PRJEB25283) (Cwiklinski et al., 2015a) using miRDeep2’s “mapper” function with default parameters. We supplied miRDeep2 with a non-redundant fasta file of the 89 *F. hepatica* miRNAs known prior to this study, gathered from published reports (Xu et al., 2012b; Fontenla et al., 2015a; Fromm et al., 2015b; Ovchinnikov et al., 2020). We ran miRDeep2 on all libraries individually, then combined the six library datasets for miRNA calling. Within this combined dataset, we accepted miRNAs that met all of the following criteria: (i) ≥10 reads mapping to the mature sequence; (ii) ≥1 read mapping to a star/passenger sequence; (iii) A precursor predicted to fold into a stable hairpin, supported by a significant randfold p-value. Some partner sequences were retained without passing all these filters, because their opposite strand partner miRNA did pass muster. We classified 5p and 3p variants according to their relative locations on the precursor RNA.

### 3.3 Developmental expression qPCR

Following RNA extraction from three biological replicates of each sample type, comprising 200 metacercariae, 200 NEJs, single adult parasites, or EV batches collected as described above, 5 ng of each RNA was reverse transcribed to cDNA using the miRCURY reverse transcription kit (Qiagen). Qiagen’s miRCURY Locked Nucleic Acid (LNA) assays were designed for every miRNA in our version of the *F. hepatica* miRNome. These assays were run against cDNA replicates with the miRCURY SYBR Green Mastermix (Qiagen) on a Qiagen RotorGene Q qPCR instrument. For data analysis, Ct data were extracted using the RotorGene software suite. Targets were considered “expressed” where they were amplified in at least two of the three replicates. Fold change analysis used the ΔΔCt method (Pfaffl, 2001), employing the dataset mean Ct (30.23±1.57) as a reference value. Negative controls replaced cDNA with water. Some LNA assays exhibited repeated unresolvable non-specific amplification in negative controls, these have been omitted from the dataset.

### 3.4 miRNA target prediction, correlation, functional and network analysis

To identify potential mRNA targets for *F. hepatica* miRNAs we adapted a consensus prediction method as described by (Gillan et al., 2017). After gathering 3’UTR sequences from *Bos taurus* (Ensembl version 104 (Howe et al., 2021)) and *F. hepatica* (WormBase ParaSite project PRJEB25283 version WBPS16 (Cwiklinski et al., 2015a; Bolt et al., 2018)), we used three miRNA target prediction algorithms (miRANDA (Enright et al., 2003), PITA (Kertesz et al., 2007) and RNAhybrid (Rehmsmeier et al., 2004)) to match all *F. hepatica* miRNAs with *F. hepatica* UTRs, and adult EV miRNAs with *B. taurus* UTRs. In both cases we took a consensus approach, using a custom Python script to identify and retain only those mRNA:miRNA pairs that were identified by all three algorithms. Within each tool, we used the same thresholds as (Gillan et al., 2017): miRanda, total score >145, energy < −10; RNAhybrid, p<0.1, energy < −22; PITA, seed sequence of 8 bases with ΔΔG < −10.

For endogenous miRNA targets in *F. hepatica*, we gathered TPM expression data for target transcripts from the *F. hepatica* transcriptome (hosted on WormBase ParaSite (Cwiklinski et al., 2018; Cwiklinski et al., 2021)), and generated correlation coefficients (CC) for each miRNA (qPCR Ct) and mRNA (RNA-Seq TPM) pair, using Excel’s “CORREL” function. We accepted only those pairs with CC>0.950 for further analysis. Each mRNA was manually annotated by BLASTx against the ncbi nr dataset using DIAMOND BLASTx (Buchfink et al., 2015), where the top hit scoring e<0.001 was accepted. For miRNA targets in ruminant hosts, hit pairs were filtered against matching transcripts shown to be down-regulated in lymph node (Naranjo-Lucena et al., 2021), peripheral blood mononuclear cell (PBMC) (Alvarez Rojas et al., 2016; Fu et al., 2016; Garcia-Campos et al., 2019) or liver tissue transcriptomes (Alvarez Rojas et al., 2015), during sheep or cow *F. hepatica* infections. Remaining mRNAs were converted to a list of human gene names via UniProt (UniProt, 2021), and then searched for pathway analysis using Reactome (Jassal et al., 2020). Networks were generated in Cytoscape (Shannon et al., 2003), with editing for colour, image size and text clarity in Inkscape (https://inkscape.org/).

## 4 RESULTS

### 4.1 Redundancy and diversity within published *F. hepatica* miRNA datasets

An initial manual analysis of all published *F. hepatica* miRNAs (including the eight *F. gigantica* miRNAs reported by (Xu et al., 2012a) yielded a redundant total of 186 mature miRNA sequences across four publications (Supplementary Table 1) (Xu et al., 2012b; Fontenla et al., 2015a; Fromm et al., 2015b; Ovchinnikov et al., 2020). Naming of miRNA sequences was inconsistent between individual papers and with the originally described miRBase naming system (Ambros et al., 2003), necessitating manual analysis and clustering to generate a non-redundant list before beginning our analysis. Where sequence variants of different lengths had been reported, we retained the longest version in our non-redundant list. We established the previously published miRNome at 89 non-redundant mature *F. hepatica* miRNA sequences, plus eight *F. gigantica* miRNAs which we retained for qPCR analysis (see section 4.4 and Supplementary Table 1). Sequence variants of individual miRNAs were visible between publications but for the most part, variation was at the miRNA 3’ end, with seed regions remaining intact and consistent. Each non-redundant miRNA (Supplementary Table 1) was given a consensus name according to sequence similarity with known miRNAs, for ease of notation throughout this paper (Supplementary Table 1 also notes the original author designated names for comparison). Previously published *F. hepatica*-specific miRNAs were named “fhe-pubnovel-*n*” and arbitrarily numbered. Note that the total of 89 sequences considers 5p and 3p sequences from the same hairpin as separate mature sequences. Note that after we completed this part of the study in 2020, a similar analysis was published by other authors (Ricafrente et al., 2020).

### 4.2 Discovery of miRNAs across *F. hepatica* developmental stages

For *de novo* discovery of expressed miRNAs we performed small RNA sequencing of multiple *F. hepatica* life stage libraries (metacercariae, non-growing NEJ, growing NEJ, adult, and EVs from both non-growing NEJ and adult). These datasets were combined and then qualitatively analysed for miRNAs using miRDeep2. This approach yielded 91 mature miRNA sequences, 29 of which had been previously reported in *F. hepatica* (Supplementary Table 2). Most sequences (61) were newly described miRNAs that lacked matches in miRBase or miRGeneDB searches, and therefore appear to be unique to *F. hepatica*. These data expand the known miRNA complement of *F. hepatica* by 40%, extending the *F. hepatica* miRNome to 150 sequences. Supplementary Datasheet 1 details this updated version of the *F. hepatica* miRNome. The absence of biological replicates prevented RNA-Seq based differential expression analysis across our datasets. However, presence/absence analysis did highlight differences in miRNA complements between libraries. Figure 1 shows that of the 91 miRNA orthologues detected, all apart from six were found in multiple libraries, with four (fhe-mir-125a-5p, fhe-mir-1989-5p, fhe-mir-277-3p and fhe-mir-71b-5p) present in all six libraries. Each library contained 27 miRNA orthologues apart from NEJ EVs, which contained 12. Library-specific miRNAs were seen in metacercariae (fhe-mir-2b-2-5p), NEJ (fhe-mir-31-5p, fhe-pubnovelmir-4-3p), adult (fhe-pubnovelmir-23-5p, fhe-pubnovelmir-4-5p) and NEJ_EV (fhe-mir-125b-5p, fhe-mir-2c-5p) samples. All miRNAs were subsequently assayed using qPCR, as described in section 4.4.

**Figure 1.**
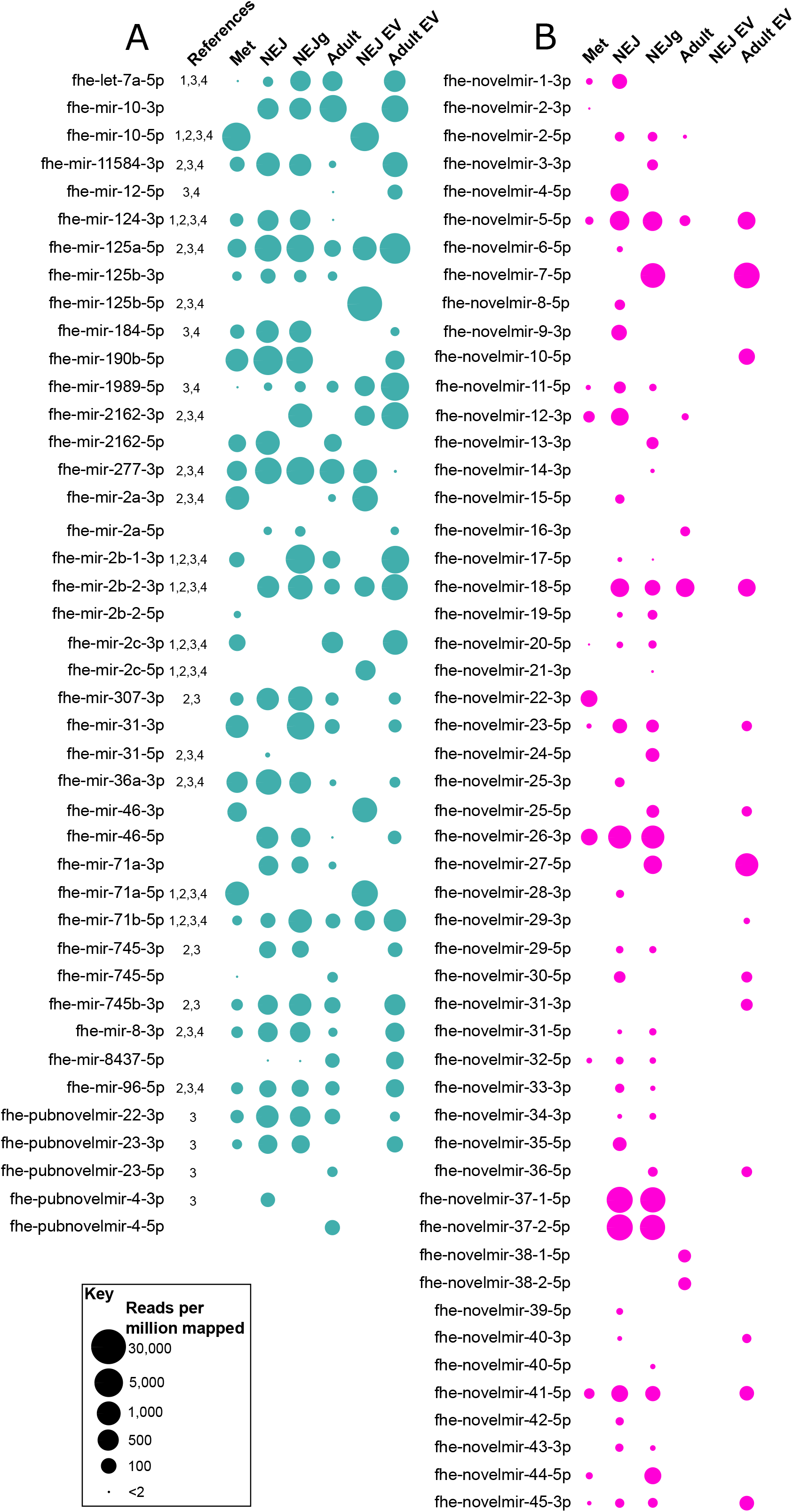
Presence/absence of novel micro (mi)RNAs across *Fasciola hepatica* RNA-Seq libraries. Presence of miRNA is indicated by a bubble, with bubble diameter representative of mature miRNA read coverage normalized per million reads mapped to the F. hepatica genome (see scale). Panel A describes previously published miRNAs, panel B shows novel miRNAs. Libraries are metacercariae (Met), newly excysted juvenile (NEJ), NEJ growing in presence of serum (NEJg), adult ex-vivo parasites (Adult), extracellular vesicles (EV) from NEJ (NEJ EV) and EV from adult (Adult EV). All datapoints represent one biological replicate. Names are consensus titles, see Supplementary Table 1 for previously used naming; see Supplementary Table 2 for complete data. References: 1,(Xu et al., 2012a); 2,(Fontenla et al., 2015b); 3,(Fromm et al., 2015c); 4,(Ovchinnikov et al., 2020).

Our sequencing yielded the first opposite strand versions of seven previously reported miRNAs (fhe-mir-10, fhe-mir-125b, fhe-mir-2a, fhe-mir-2162, fhe-mir-31, fhe-mir-71a, fhe-mir-745, fhe-pubnovelmir-4; Figure 1, Figure 2), and both 5p and 3p variants of the previously unreported fhe-mir-46. Our dataset also confirmed the previously reported *F. hepatica*-specific miRNAs fhe-pubnovelmir-4, -22 and -23 (Fromm et al., 2015c). These data provide further confirmation of these sequences as being *bona fide* miRNAs, and suggests that miRNA 5p and 3p variants may be differentially regulated across fluke life cycle transitions.

**Figure 2.**
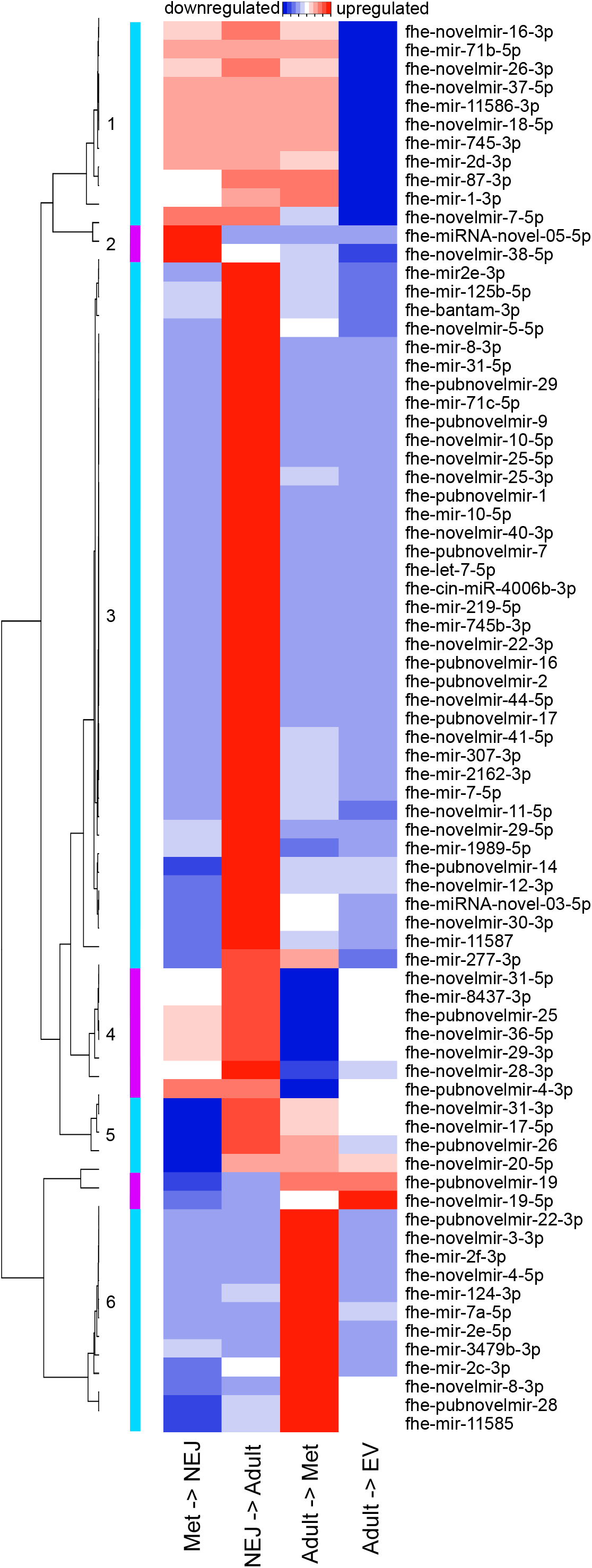
Differential expression of micro (mi)RNAs across *Fasciola hepatica* life stage transitions. Heatmap displays mean fold change in miRNA expression, as detected by quantitative (q)PCR, between the indicated life stage libraries. Blue indicates downregulation, red indicates upregulation, white indicates no change. Six expression profiles are indicated by cyan and magenta bars numbered 1-6.

### 4.3 Novel *F. hepatica* miRNAs

In addition to the previously reported miRNAs described above, our sequencing also yielded 52 previously unrecognised and currently *F. hepatica* specific miRNAs (fhe-novelmir-1 through -45; Supplementary Table 2). These novel sequences were much less abundant across libraries (mean 80 RPMM) than the previously reported sequences (mean 593 RPMM). Further, novel miRNAs were more variable in presence/absence than known miRNAs (Figure 1), with thirteen in the metacercaria library, around 30 in NEJ, seven in adult, 14 in adult EV, and none in NEJ EV. Stage-specific miRNAs were found in metacercariae (fhe-novelmir-2-3p, fhe-novelmir-22-3p), non-growing NEJ (fhe-novelmir-4-5p, fhe-novelmir-6-5p, fhe-novelmir-15-5p, fhe-novelmir-25-3p, fhe-novelmir-28-3p, fhe-novelmir-35-5p, fhe-novelmir-39-5p, fhe-novelmir-42-5p), growing NEJ (fhe-novelmir-3-3p, fhe-novelmir-13-3p, fhe-novelmir-14-3p, fhe-novelmir-21-3p, fhe-novelmir-24-5p, fhe-novelmir-40-5p) adult (fhe-novelmir-16-3p, fhe-novelmir-38) and adult EV (fhe-novelmir-10-5p, fhe-novelmir-29-3p, fhe-novelmir-31-3p).

### 4.4 Profiling miRNA developmental expression across life stage transitions

To quantify the regulation of miRNA expression across life stage transitions, we purchased LNA-based qPCR assays for each of the 150 miRNAs in our version of the *F. hepatica* miRNome, plus the *F. gigantica* miRNAs reported by (Xu et al., 2012a) (Supplementary Datasheet 1). In assays covering three biological replicates each of met, NEJ, adult, and adult EV libraries, 76 miRNAs were detected in at least one sample type (Supplementary Table 3). Twenty miRNAs were detected in a single life stage, with 18 miRNAs detected in all four libraries. Differential expression analysis identified up to six distinct expression profiles (Figure 2), demonstrating that the majority of miRNAs were expressed most highly in metacercariae, showing downregulation during transition to NEJ, and then upregulated during transition from NEJ to adult. In addition to tissue samples, we also analysed adult derived EVs, in which 28 miRNAs were detected. Following normalisation, most of these were less abundant than in adult tissue, but two were relatively more abundant in EV than tissue samples (fhe-pubnovelmir-19, 179 fold upregulated; fhe-miRNA-novel-06-3p, 549 fold upregulated). The latter miRNA is of *F. gigantica* origin (Xu et al., 2012a); we also amplified an additional three *F. gigantica* miRNAs (fhe-miRNA-novel-03-3p, fhe-miRNA-novel-05-3p, fhe-miRNA-novel-09-3p), suggesting that orthologues of these are expressed by *F. hepatica*. NEJ-derived EVs were generated and assayed for miRNA expression, but these proved challenging to detect by qPCR; these data have not been included here.

### 4.5 Predicting mRNA targets for cellular and secreted miRNAs

*F. hepatica* EVs can be internalised by host cells ((Marcilla et al., 2012; de la Torre-Escudero et al., 2019) and seemingly release miRNA cargo with potential to regulate host mRNA expression (Tran et al., 2021). To explore miRNA:mRNA regulatory networks, we first used target prediction algorithms to identify interactions between the 28 miRNAs detected by qPCR in adult EVs (Figure 2) and host (bovine) mRNA 3’UTRs. All 28 adult EV miRNAs matched with one or more of 2281 cow mRNAs. Given that one of the primary effects of miRNA binding is to reduce stability and expression of target mRNAs, we then filtered these target genes to identify which were downregulated during *F. hepatica* infection. We achieved this by mining of published transcriptomes from sheep lymph node (Naranjo-Lucena et al., 2021), sheep and cow PBMCs from both acute and chronic fasciolosis scenarios (Alvarez Rojas et al., 2016; Fu et al., 2016; Garcia-Campos et al., 2019) and sheep liver tissue (Alvarez Rojas et al., 2015), all recovered from experimental *F. hepatica* infections in comparison with time matched uninfected controls. These comparisons identified 298 miRNA-targeted downregulated transcripts in lymph node, 57 targets from acute infection PBMCs from sheep, 20 targets from chronic infection PBMCs from sheep, 78 targets from chronic infection PBMCs from cow, and three targets from sheep liver. No matches were detected with transcripts downregulated in PBMCs from cow acute infections. Of the 397 total target mRNAs, Reactome analysis identified 124 pathways within 23 top-level pathway families (Figure 3). Amongst the latter, the largest proportion of transcripts (23%) mapped to Signal Transduction, with Immune System (16%), Metabolism (11%), Metabolism of Proteins (7%), Transport of Small Molecules (7%), Gene Expression (Transcription) (6%) and Extracellular Matrix Organisation (4%) also represented in the top 75% of target transcripts. The remaining 25% incorporated an additional 16 top level pathways, detailed in Figure 3 and Supplementary Table 4. Overrepresentation analysis identified 68 pathways scoring p<0.05, with the most statistically significant target pathways being FOXO-mediated transcription of cell cycle genes (p=1.24E-04), and Interleukin-4 and Interleukin-13 signalling (p=3.96E-04) (Supplementary Table 5). Of the secreted *F. hepatica* miRNAs, fhe-mir-745b-3p matched the largest number of bovine mRNAs (365), followed by fhe-miRNA-novel-03-5p (361), fhe-pubnovelmir-7 (306) and fhe-pubnovelmir-17 (210).

**Figure 3.**
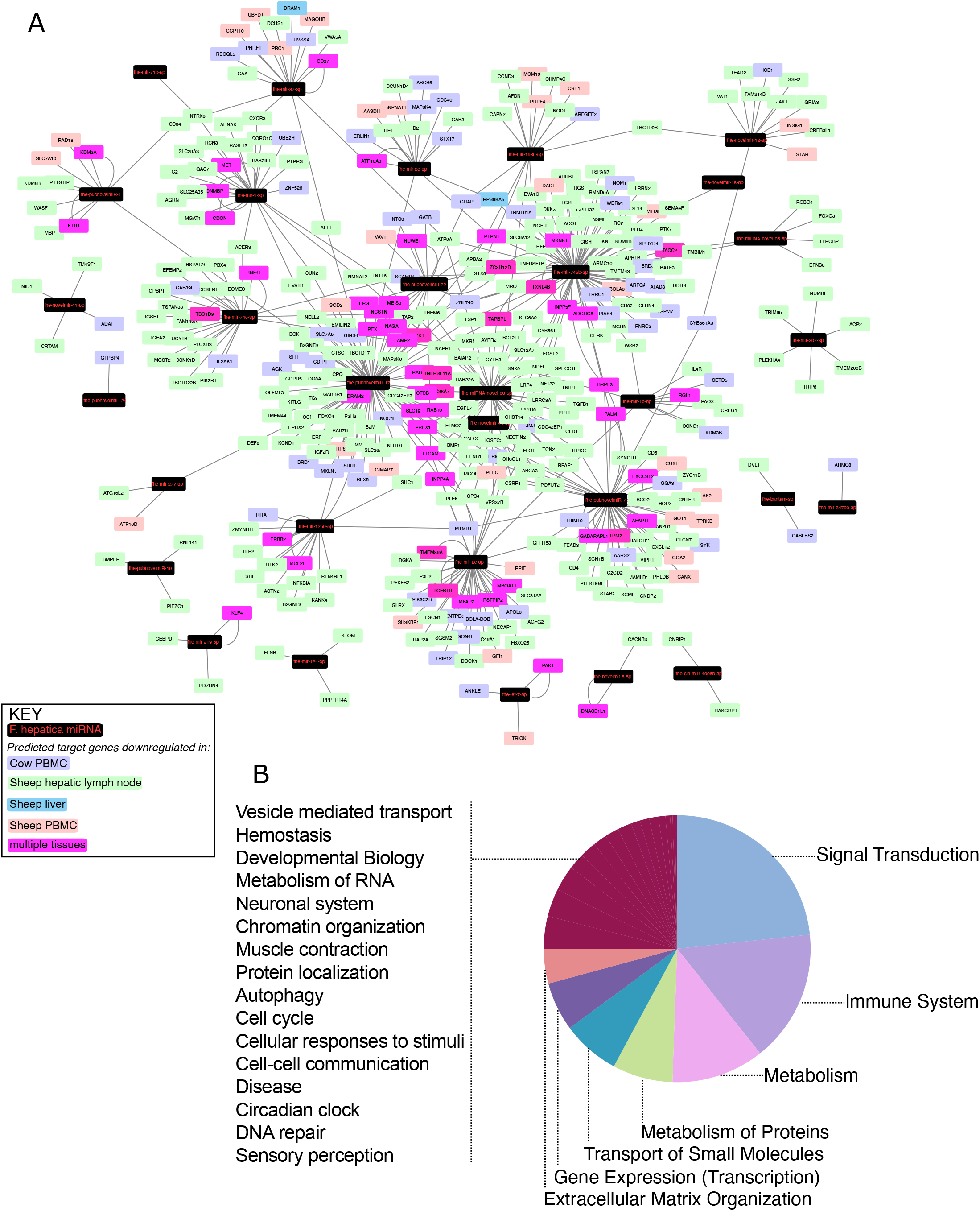
Network analysis of bovine (*Bos taurus*) mRNAs downregulated in fasciolosis and targeted by micro (mi)RNAs secreted in Extracellular Vesicles of adult *Fasciola hepatica*. (A) Nodes represent miRNAs (black rectangles) or bovine mRNA targets (ovals, coloured according to originating transcriptome dataset as detailed in Key). (B) Reactome pathway membership of downregulated bovine transcripts classified by top level Reactome pathway. For full analysis data, see Supplementary Tables 4 and 5.

To uncover possible regulators of fluke gene expression during intra-mammalian development, we next investigated interactions between the tissue-expressed miRNome and the endogenous *F. hepatica* mRNA transcriptome (Cwiklinski et al., 2018; Cwiklinski et al., 2021), identifying 3837 miRNA:mRNA pairs, within which 155 miRNAs matched 2307 mRNA 3’UTR targets. These target transcripts contained a wide range of biological targets and functions. We filtered this large number of pairs using expression correlation analysis of each of the 78 miRNAs detected in met, NEJ or adult tissue by qPCR (Ct number), with their matching mRNAs (RNA_Seq TPM) across met, NEJ and adult tissues. This highlighted 296 inversely correlated miRNA:mRNA pairs with a correlation coefficient (CC) >0.950 (Supplementary Table 6; Figure 4). The largest proportion of these targets (*n*=97; 24 %) were transcripts encoding hypothetical or unannotated proteins with unknown homology or function, which therefore evaded functional annotation. The remainder included transcripts related to transcriptional and translational control, metabolic enzymes, structural and cytoskeletal components, signal transduction and proteolysis. Of particular interest was the identification of putative miRNA interactions with key targets that have been noted in *Fasciola* literature as biologically important, and/or appealing targets for control. These included neuromuscular transcripts (voltage/ligand gated ion channels, transient receptor potential channels, G protein-coupled receptors, signal peptidase components, neuropeptide receptors, heterotrimeric G protein components, calmodulins), secreted metabolic modulators and nutrient scavengers (fatty acid binding protein, ferritin, glutathione transferase) secreted proteases (cathepsin L, legumain, cercarial protease), and individual components of exosome and glycan biosynthesis.

**Figure 4.**
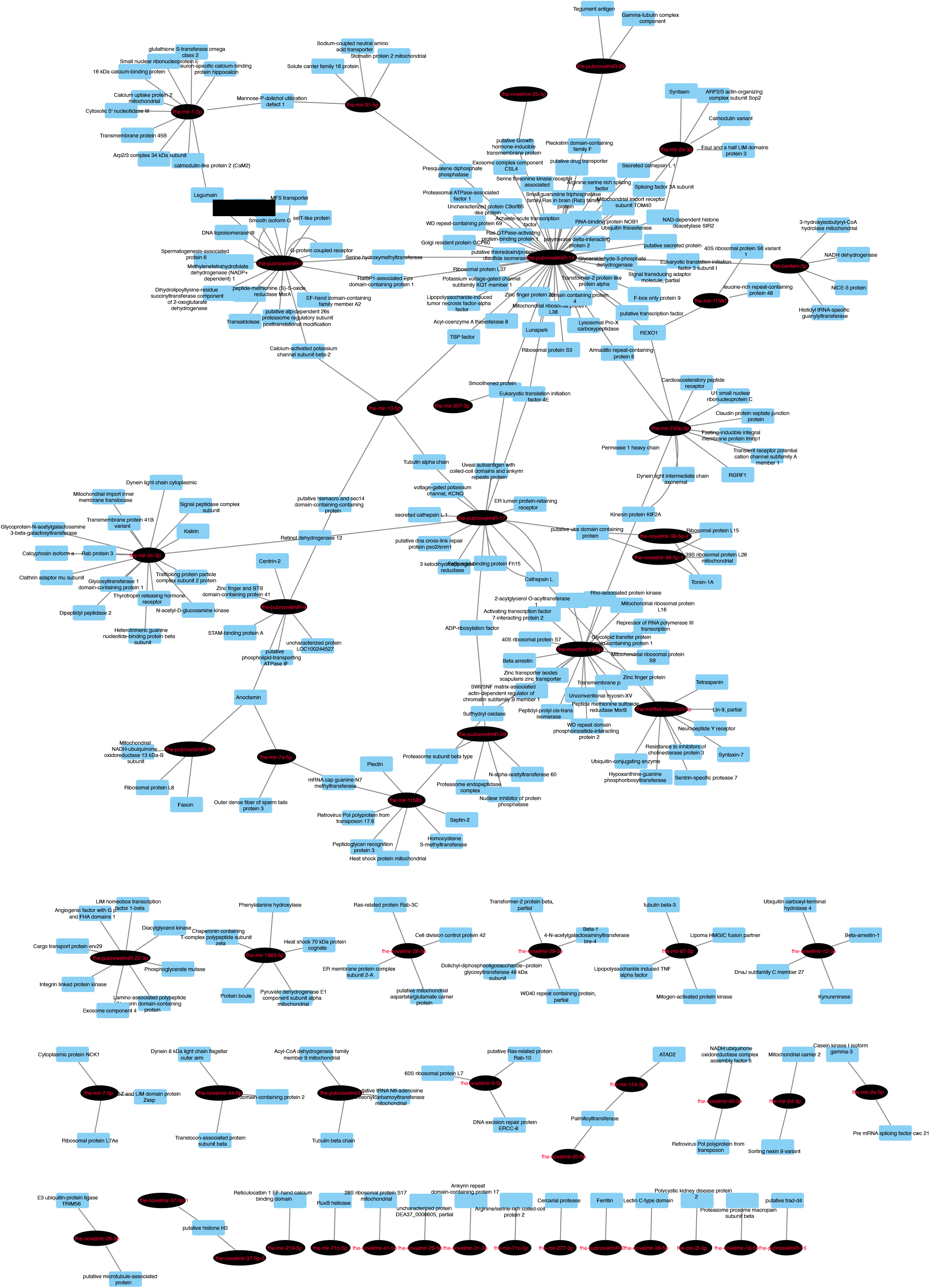
(A) Network analysis of predicted endogenous *Fasciola hepatica* mRNA:miRNA interactions. Nodes represent miRNAs (black ovals) or mRNA targets (blue rectangles). Target transcripts encoding hypothetical proteins or proteins lacking database homology have been omitted. For full analysis data, see Supplementary Table 6.

## 5 DISCUSSION

*F. hepatica* remains an under-studied pathogen, at least relative to other flatworm parasites such as *Schistosoma* spp. blood fluke. Ongoing difficulties with parasite control necessitate new control approaches; a clear way to address this need is through improved understanding of fundamental parasite biology and host-parasite interactions. This improved knowledge can lead to identification of new diagnostic and control opportunities (Cwiklinski and Dalton, 2018; McVeigh et al., 2018). This study contributes to understanding of the *F. hepatica* miRNome, which prior to this work was represented by six studies that have identified and begun functional classification of liver fluke miRNAs (Xu et al., 2012b; Fontenla et al., 2015a; Fromm et al., 2015b; Ovchinnikov et al., 2020; Ricafrente et al., 2020; Tran et al., 2021). The data presented in this paper have extended the known *F. hepatica* miRNome to 150 mature miRNAs, provided the first developmental profile of miRNAs in intra-mammalian *F. hepatica* life stages, and attached comprehensive computational predictions of miRNA-mRNA functional regulatory networks for both cellular and secreted miRNAs.

Prior to data collection, this project began with a manual analysis of all published *F. hepatica* miRNAs, yielding 186 mature miRNA sequences reported across four publications (Supplementary Table 1) (Xu et al., 2012b; Fontenla et al., 2015a; Fromm et al., 2015b; Ovchinnikov et al., 2020). Many sequences were replicated or named differently between papers, or named in a fashion that was inconsistent with sequence orthology, and/or with the seminal miRBase naming system (Ambros et al., 2003). This is a confusing situation for researchers attempting to understand the *F. hepatica* miRNome. Recently, other authors have recognised this issue, addressing it with updated (but non-miRBase) naming terminology (Ovchinnikov et al., 2020) and re-analysing the miRNome *in toto* (Ricafrente et al., 2020). As part of our re-analysis, we assigned consensus names to all published miRNAs in line with the miRBase naming system (Ambros et al., 2003). Our primary goal in this was to generate a consensus dataset for our own reference, but we present it in Supplementary Table 1 as a dataset that may be useful for others. Our manual clustering of the published *F. hepatica* miRNA complement yielded 89 distinct mature miRNA sequences (Supplementary Table 1). This is a greater number than the 77 reported by (Ricafrente et al., 2020), who omitted deemed some miRNAs from their final total, deeming them invalid. We retained all previously reported sequences in our non-redundant list, since one of our goals was to test the existence of these sequences with qPCR assays.

Our initial miRNA sequencing analyses yielded qualitative identification of 91 miRNAs across multiple *F. hepatica* life stages, representing the largest single miRNA dataset reported for *F. hepatica*. This total comprised 29 previously reported miRNAs and 61 novel, and apparently *Fasciola*-specific miRNAs (we consider these sequences novel since they did not match any held by miRBase or miRGeneDB, and were not found within supplementary data files of previously published flatworm miRNA papers). These therefore can currently be considered *F. hepatica*-specific miRNAs. Discovery of so many new sequences in one dataset is not unprecedented (Winter et al., 2012), and may suggest the rapid evolutionary rates of miRNA genes (Britton et al., 2020) within Fasciolidae. This hypothesis could be confirmed by discovery of orthologues of our 61 novel sequences in *F. gigantica* and other closely related flukes.

The single RNA-Seq biological replicates generated here were used only for qualitative miRNA discovery; the absence of biological replicates made them inappropriate for quantitative analyses and therefore did not permit differential expression analysis across our datasets. We addressed this by instead using qPCR to separately measure expression of miRNome components across life stages. Seventy-six miRNAs (including four orthologues of presumed *F. gigantica*-specific miRNAs (Xu et al., 2012a)) were detected in at least one life stage. Ten miRNAs were detected only in metacercariae, and ten only in adults, perhaps suggesting specific biological functions in these life stages. Overall, the majority of miRNAs were expressed most highly in metacercariae. This may be related to the induction of stasis in this life stage, where miRNAs could be involved in pausing transcription from pre-transcribed mRNAs until a host is encountered. Previous studies have shown that metacercariae do harbour abundant transcripts (Cwiklinski et al., 2015a; Cwiklinski et al., 2018), but it has not yet been proven whether these are transcribed prior to, or during, dormancy. Further work will be required to test this hypothesis.

To identify potential functions of liver fluke miRNAs, we performed computational predictions of miRNA:mRNA interactions, based on consensus matches across three different prediction algorithms. Twenty-eight miRNAs were detected in adult derived EVs. While all of these were also found in other libraries, their presence in EVs suggests their secretion by adult parasites and suggests that they may have functions in host-parasite interactions. We used three miRNA target prediction algorithms to identify mRNA 3’UTR targets from the bovine transcriptome with which secreted miRNAs could bind. To improve confidence in our predictions, we purposely selected three tools that each rely on distinct analytical methods (Peterson et al., 2014), and took a consensus approach, in which we accepted only those hits that were identified by all three tools. Resulting hits were then filtered against transcriptomes from sheep/cattle tissues to identify target transcripts downregulated during fasciolosis (since downregulation could indicate the targeting and transcriptional destruction of these transcripts by secreted fluke miRNAs associated with Argonaut complexes). These comparisons identified 298 miRNA-targeted, significantly downregulated transcripts across fasciolosis transcriptomes from lymph node, PBMCs and liver from sheep and/or cattle. That these matches were seen across these distinct tissues, and from both acute and chronic infections in sheep, suggests the wide-ranging and systemic importance that fluke secreted miRNAs might have for fasciolosis pathology and virulence. While these correlations do not prove *in vivo* interactions, they do provide a set of hypotheses for future testing. Network visualisation of these interactions (Figure 4) showed striking differentiation between miRNAs forming hub nodes, each targeting multiple host mRNAs, and nine miRNAs which formed isolated nodes, which targeted only a handful of specific mRNAs and were not part of the wider network. Functional genomics experiments will be required to test and validate the functional implications of these predictions for host-parasite interactions. Reactome pathway analysis of bovine mRNA targets showed that the majority were implicated in signal transduction or gene expression pathways, and as such could conceivably be involved in fluke-mediated immunosuppression or cellular pathology. Previous publications have also suggested roles for fluke secreted miRNAs in immunoregulation of the host (Fromm et al., 2015a; Fromm et al., 2017; Ovchinnikov et al., 2020), and used miRNA target prediction algorithms to annotate host mRNAs potentially targeted by fluke secreted miRNAs. (Ovchinnikov et al., 2020) focused on 24 *F. hepatica* adult and EV derived miRNAs (alongside 22 *S. mansoni* miRNAs), using PITA and TargetSCAN to identify 321 targeted mRNAs, each of which was predicted by both algorithms and conserved in both bovine and human hosts. These included 11 targets within WNT signalling pathways, and 23 immune system mRNAs. (Ricafrente et al., 2020) analysed interactions between 38 *F. hepatica* NEJ miRNAs (Fontenla et al., 2015b) and innate immune cell transcripts using a single algorithm (miRDB’s MirTarget tool). This identified 26 target genes expressed in eosinophils, dendritic cells and neutrophils. Both of these studies employed distinct methods to each other, and to our study, and all three studies have focused on distinct but overlapping miRNA datasets. Our approach is unique because our predictions focus on mRNA targets that are demonstrably downregulated during fasciolosis. Our data suggest that these correlations are the result of modulation of the host transcriptome by secreted fluke miRNAs and experimental verification of these molecular interactions could provide new insights into parasite-to-host communication.

Amongst the fluke miRNAs within our dataset was fhe-mir-125b-5p, which is notable for being the first helminth secreted miRNA to have an associated host-interacting function. Human hsa-mir-125b controls macrophage activation (Duroux-Richard et al., 2016). In a compelling example of convergent evolution, *Schistosoma* spp mir-125b-5p similarly triggers a pro-inflammatory phenotype in mouse macrophages (Liu et al., 2019); *F. hepatica* mir-125b-5p is expressed in NEJ parasites (Fontenla et al., 2015b), can be found associated with mammalian argonaut (Ago-2) within the peritoneal macrophages of infected mice, and is predicted to target innate immune components (Tran et al., 2021). Our analysis showed that fhe-mir-125b-5p was also expressed in metacercariae and adult parasites, and secreted in adult EVs. Reflecting this wider expression profile, our analysis of fhe-mir-125b-5p targets in host transcriptomes identified 18 genes, within reactome pathways including signaling by interleukins, receptor tyrosine kinases and NTRK3, and O-linked glycosylation of mucins (Supplementary Table 4). TRAF6, the target of fhe-mir-125b identified by (Tran et al., 2021), was not amongst our targets.

Our next goal was to investigate intracellular interactions between *F. hepatica* miRNAs and endogenous cellular fluke transcripts. We exploited publicly available transcriptomes from metacercariae, NEJ and adult fluke (Cwiklinski et al., 2018), with which we matched our qPCR expression data for 76 miRNAs detected in these same life stages. Interacting miRNA:mRNA pairs predicted by the same consensus approach as detailed above, were filtered to include only those with highly correlated inverse expression (CC>0.950). Our hypothesis was that the resulting 397 pairs of computationally matched, co-expressed miRNA:mRNA pairs would be those most likely to have a biologically relevant interaction *in vivo*. The resulting analyses (FIGURE 4, Supplementary Table 6) highlighted miRNAs correlated with, and potentially controlling expression of, a wide range of protein coding transcripts, across a range of key biological functions. Of note were links with transcripts encoding proteins of key interest for fluke control (cathepsin, glutathione transferase, nerve/muscle transcripts). Knowledge of the regulatory mechanisms controlling expression of these key targets across fluke development could open new avenues to probing and therapeutically inhibiting their functions. For example, miRNA mimics or inhibitors could be used to manipulate transcript expression alongside or in place of RNA interference (RNAi) methods. These new approaches could be useful for laboratory-based functional genomics, or even as new avenues for control (miRNAs have been suggested as therapeutic targets in various human diseases (Marracino et al., 2021; Shi et al., 2021; Smit-McBride and Morse, 2021)). Furthermore, inhibiting EV release from helminths would conceivably prevent delivery of miRNAs (and other bioactive molecules) to host immune cells. Recently, (Bennett et al., 2020) showed that a chemical inhibitor of EV biogenesis blocked the secretion of EVs from *F. hepatica in vitro* as initial proof-of-concept for such an approach.

One of the major foci for parasite derived miRNAs has been their potential use as diagnostic biomarkers (Mu et al., 2021) We have identified at least 28 EV-derived secreted miRNAs from adult parasites, which could potentially represent PCR detectable indicators of mature fluke infections. However, further experiments are needed to determine if these markers are detectable in blood or other biofluids from infected animals. At the present time this has not been demonstrated for *F. hepatica*, although small RNA sequencing has been used to detect four *F. gigantica* miRNAs in sera from infected buffalo (Guo and Guo, 2019), which provides encouraging support for the technical feasibility of this approach for *F. hepatica*.

This work has considerably expanded the known extent of the *F. hepatica* miRNome, profiled the developmental expression of miRNAs across intra-mammalian life stages, and attached functional annotations to cellular miRNAs relative to endogenous *F. hepatica* transcripts, and to EV miRNAs relative to host transcripts. These data provide new perspectives on *F. hepatica* miRNA biology and a set of testable hypotheses for future research. This dataset therefore represents an important contribution to our understanding of miRNA functions in *F. hepatica* biology, virulence and pathogenicity, and provides potential new avenues towards fasciolosis control and diagnostics.

## Supporting information

Supplementary Datasheet 1

## ACKNOWLEDGEMENTS

This work was supported by a Postgraduate Research Studentship from the Northern Ireland Department for the Economy (DFE) to C.M.H., grant BB/L019612/1 from the Biotechnology and Biological Sciences Research Council (BBSRC) to M.W.R., BBSRC grants BB/K009583/1 and BB/H009477/1 to A.G.M., and Bill and Melinda Gates Foundation Grand Challenges Explorations Grant OPP1083083 to P.M. The funders had no role in the study design or collection, analysis, and interpretation of the results.

